# Savings and interference following learning to reach with mirror reversed feedback

**DOI:** 10.64898/2026.07.28.741240

**Authors:** Sarvenaz Heirani Moghaddam, Gerome A. Manson, Erin K. Cressman

## Abstract

Learning to reach with a visuomotor distortion has been shown to influence subsequent reaches with the same distortion and with a new distortion. Here, we examined whether learning to reach with a small 20° mirror reversed distortion leads to faster re-learning of the same distortion (i.e., demonstrates savings) and whether learning to reach with a mirror reversed distortion influences subsequent reaches with a 20° visuomotor rotation distortion. Thirty participants first learned to reach with the mirror reversed distortion. Following washout trials with aligned cursor feedback, 15 participants reached again with the mirror reversed distortion (MR–MR group), while the 15 other participants reached with a visuomotor rotation distortion (MR–VMR group). An additional twenty participants only reached with the visuomotor rotation distortion (VMR-only group). Implicit (unconscious) and explicit (conscious strategy) contributions to learning were assessed using the process dissociation procedure. Evidence of savings was evident in the MR–MR group, such that participants demonstrated reduced hand angles when re-introduced to the mirror reversed distortion. This savings was driven by explicit processes, consistent with the rapid retrieval of previously acquired task solutions. Additionally, learning to reach with the mirror reversed distortion interfered with learning to reach with the visuomotor rotation distortion, such that the MR-VMR group demonstrated increased reach variability and longer reaction times when reaching with the visuomotor rotation distortion compared to the VMR-only group. Reduced implicit contributions were also evident in the MR-VMR group compared to the VMR-only group. Together, results indicate that learning to reach with a mirror reversed distortion promotes savings and influences learning to reach with a visuomotor rotation distortion through engagement of explicit processes.

## Introduction

Learning to reach with a novel visuomotor mapping has been shown to shape later reaching performance. The impact of learning to reach with a visuomotor distortion on later learning has been examined in the lab using visuomotor rotation paradigms, in which visual feedback of the hand’s trajectory is rotated clockwise (CW) or counterclockwise (CCW) by a fixed angle (e.g., 20°) relative to actual hand motion (Baraduc & Wolpert, 2002; Bastian, 2008; Ghahramani et al., 1996; Vetter et al., 1999). In visuomotor rotation paradigms, re-learning the same cursor rotation demonstrates savings, characterized by faster error reduction when the cursor rotation is re-introduced a second time relative to initial learning (de Brouwer et al., 2018; Morehead et al., 2015). Facilitation of learning has also been observed when the second cursor rotation is in the same direction as the initially learned cursor rotation, even if the magnitude of the distortion differs (e.g., a 20° CW cursor rotation followed by a larger 40° CW cursor rotation). In contrast, interference of learning is evident when the second rotation is in the opposite direction to the initial rotation (e.g., 20° CW cursor rotation followed by a 20° CCW cursor rotation; Werner et al., 2015; Brashers-Krug et al., 1996; Krakauer et al., 2005). Savings and facilitation are often attributed to the re-engagement of previously established learning processes, whereas interference may arise when previously engaged processes are inappropriate for the new mapping and must be suppressed or reconfigured (Bond & Taylor, 2015; Desrochers et al., 2020; Krakauer et al., 2005).

Learning processes include implicit (i.e., unconscious; Izawa et al., 2012; Izawa & Shadmehr, 2011; Shadmehr et al., 2010) and explicit processes (i.e., conscious reach strategies; Benson et al., 2011; McDougle & Taylor, 2019; Taylor et al., 2014; Werner et al., 2015). In general, the relative engagement of implicit and explicit processes during initial learning vary as a function of cursor distortion size (Modchalingam et al., 2019; Neville & Cressman, 2018; Werner et al., 2015). For small cursor rotations (e.g., cursor rotations less than 30°), initially learning to reach with the distortion is supported by implicit processes with minimal contributions from explicit processes, whereas larger rotations (e.g., cursor rotations greater than 40°) tend to engage both implicit and explicit processes (Bond & Taylor, 2015; Werner et al., 2015). Explicit processes have been strongly implicated in savings and facilitation, when previously established reach strategies can be re-engaged (Avraham et al., 2021; Bond & Taylor, 2015; Morehead et al., 2015). For example, Morehead et al. (2015) demonstrated that participants rapidly reinstated previously learned reach strategies upon re-exposure to the same visuomotor rotation distortion, resulting in reduced initial error during re-learning. In contrast, implicit processes showed little evidence of supporting savings (also see Bond & Taylor, 2015). Together, these findings suggest that the extent to which implicit and explicit processes are engaged during initial learning may determine how initial learning influences later motor learning.

In contrast to learning to reach with a visuomotor rotation distortion, learning to reach with a mirror reversed distortion has been shown to rely primarily on explicit contributions, with minimal implicit contributions even when the mirror reversed distortion is small in magnitude (Heirani Moghaddam et al., 2026; Wang & Taylor, 2021; Wilterson & Taylor, 2021). When reaching with a mirror reversed distortion, cursor feedback is mirrored across the body midline. Thus, in contrast to a visuomotor rotation distortion, the magnitude and direction of the errors experienced when reaching with a mirror reversed distortion vary across target locations.

In the current research, we asked whether explicit processes engaged during learning to reach with a small mirror reversed distortion support savings when the same distortion is re-introduced and whether these processes influence subsequent learning of a small visuomotor rotation distortion, which is primarily supported by implicit processes (Neville & Cressman, 2018; Werner et al., 2015). If re-learning depends on the re-engagement of previously established learning processes, then the explicit processes engaged during initial learning should be readily retrieved when the same mirror reversed distortion is encountered again, and savings should be evident. Moreover, these same explicit processes may interfere with learning to reach with a small visuomotor rotation distortion, as they are inappropriate for the new visuomotor mapping.

To test the influence of learning to reach with a small mirror reversed distortion on subsequent learning, participants first reached with a small mirror reversed distortion. Participants then completed a washout phase before reaching with the mirror reversed distortion again (MR-MR group) or a visuomotor rotation distortion (MR-VMR group). In addition to assessing learning to reach with the visuomotor distortions, we also quantified implicit and explicit contributions to learning using the process dissociation procedure (PDP; Jacoby, 1991; Werner et al., 2015). Within the PDP, participants completed reaches under two sets of instructions: exclusion and inclusion instructions. Under exclusion instructions, participants were asked to reach directly to the target as they did during baseline following reaches with aligned cursor feedback, without using any strategies or adjustments developed during learning trials with the visuomotor rotation or the mirror reversed distortion. Under inclusion instructions, participants were asked to use any strategies or adjustments they had learned to get the cursor to the target when the visuomotor distortion was present. Implicit learning was defined as the hand angles observed under exclusion instructions, whereas explicit learning was established based on differences in hand angles between inclusion and exclusion trials.

In accordance with previous research, we expected that explicit processes would be responsible for learning to reach with a small mirror reversed distortion, with minimal contributions from implicit processes (Heirani Moghaddam et al., 2026; Wang & Taylor, 2021; Wilterson & Taylor, 2021). We further hypothesized that savings would be observed when participants re-learned to reach with the same mirror reversed distortion. Finally, we hypothesized that learning to reach with the mirror reversed distortion would interfere with subsequent reaches with the visuomotor rotation distortion, given that the explicit contributions have been shown to persist over time (Bouchard & Cressman, 2021; Morehead et al., 2015). Clarifying the impact of learning to reach with a mirror reversed distortion on later reaches with a visuomotor rotation and mirror reversed distortion will advance our understanding of how the processes engaged during initial learning shape future motor learning.

## Methods

### Participants

Thirty participants (F = 19), aged 18–35 years (mean age = 22 ± 3.4 years), were recruited from the Queen’s University community. Participants were assigned to one of two groups: the MR–MR group (N = 15; F = 9) or the MR–VMR group (N = 15; F = 10). Participants in the MR–MR group completed two blocks of reaches with a mirror reversed distortion separated by a washout block, in which participants reached with aligned cursor feedback. Participants in the MR-VMR group were split in half, such that 7 participants reached with a clockwise (CW; 4 females) visuomotor rotation distortion and 8 participants reached with a counterclockwise (CCW; 6 females) visuomotor rotation distortion. These reaches with either the CW or CCW cursor rotation were completed after a block of reaches with the mirror reversed distortion and washout trials. In addition, we included data from a sample of 20 participants (mean age = 21 ± 3 years) who only reached with a 20° visuomotor rotation distortion. This dataset was collected previously in our laboratory and has been reported in another manuscript (Heirani Moghaddam, Manson, Cressman, submitted. Half of these participants reached with a CW cursor rotation (VMR-CW; N = 10, 7 females) and the other half reached with a CCW cursor rotation (VMR–CCW; N = 10, 7 females). Including this VMR-only group enabled us to isolate the effects of learning to reach with a mirror reversed distortion on learning to reach with a visuomotor rotation distortion in our MR-VMR group.

Only participants who demonstrated learning when initially reaching with the mirror reversed distortion were included in the analyses below. Participants in the MR-MR and MR-VMR groups were designated as non-learners based on the absence of changes in reaches with the mirror reversed distortion in the first learning block relative to the baseline block. Specifically, a non-learner designation was assigned if one of the two following criteria was met: a) mean hand angle for the late learning trials in learning block 1 with the mirror reversed distortion for either the right or left target was within 3 standard deviations of a participant’s mean late hand angle in their baseline block for the same target and/or b) reaches were in the incorrect direction. Based on these criteria, 3 participants in the MR–MR group and 2 participants in the MR–VMR group were classified as MR non-learners. Overall, the analyses below consisted of three groups: MR-MR (N = 12), MR-VMR group (N = 13), VMR-only group (N = 20).

All participants were right-handed as confirmed by the Edinburgh Handedness Inventory (mean score = 93.7 ± 8.6) and reported no history of neurological or musculoskeletal disorders. Written informed consent was obtained prior to participation. The study protocol was approved by the Queen’s University Health Sciences and Affiliated Teaching Hospitals Research Ethics Board (HSREB).

### Experimental apparatus and Procedures

The experimental protocol was similar to that used previously (see Heirani Moghaddam, Cressman, & Manson, submitted). Participants performed slicing (shooting) movements with their right hand using the Kinarm Exoskeleton (Kinarm, Kingston, ON, Canada; Kasuga et al., 2022; Scott, 1999; Singh & Scott, 2003). The Kinarm was positioned adjacent to the experimenter’s computer workstation and consisted of a downward-facing computer monitor (120 Hz refresh rate) and a reflective surface located 20.5 cm beneath the monitor. The downward-facing monitor projected visual stimuli onto the reflective surface, such that cursor feedback appeared spatially aligned with the participant’s hand, which was 20.5 cm below the reflective surface. Participants were seated in a height-adjustable wheelchair, and their arms were placed in adjustable troughs of the exoskeleton, with the left arm remaining stationary throughout the experiment. The experimenter adjusted the chair height and distance from the reflective surface to ensure comfortable viewing and reaching to the visual stimuli. Participants performed flexion, extension, abduction and adduction movements of the right arm in the horizontal plane. The right index finger’s position was displayed as a cursor (0.5 cm diameter white circle) and recorded at a sampling rate of 1000 Hz. To eliminate visual feedback of the right limb, a drape was secured around the participant’s neck with Velcro. The drape, along with the reflective surface, fully obscured the participant’s view of their right arm.

### Trials

In general, participants made slicing movements towards two targets positioned 10 cm from the home position (1 cm in diameter), which was in line with the body midline, approximately 15 cm anterior to the participant. The targets were blue circles, 0.75 cm in diameter, and appeared with equal probability. The right target was positioned 10° to the right of the y-axis, which was aligned with a participant’s midline, and the left target was positioned 10 to the left of the y-axis (Figure 1A). The *target array* was defined as the circumference of an unseen circle centered on the home position with a radius of 10 cm, corresponding to the distance between the home position and target locations.

**Figure 1.**
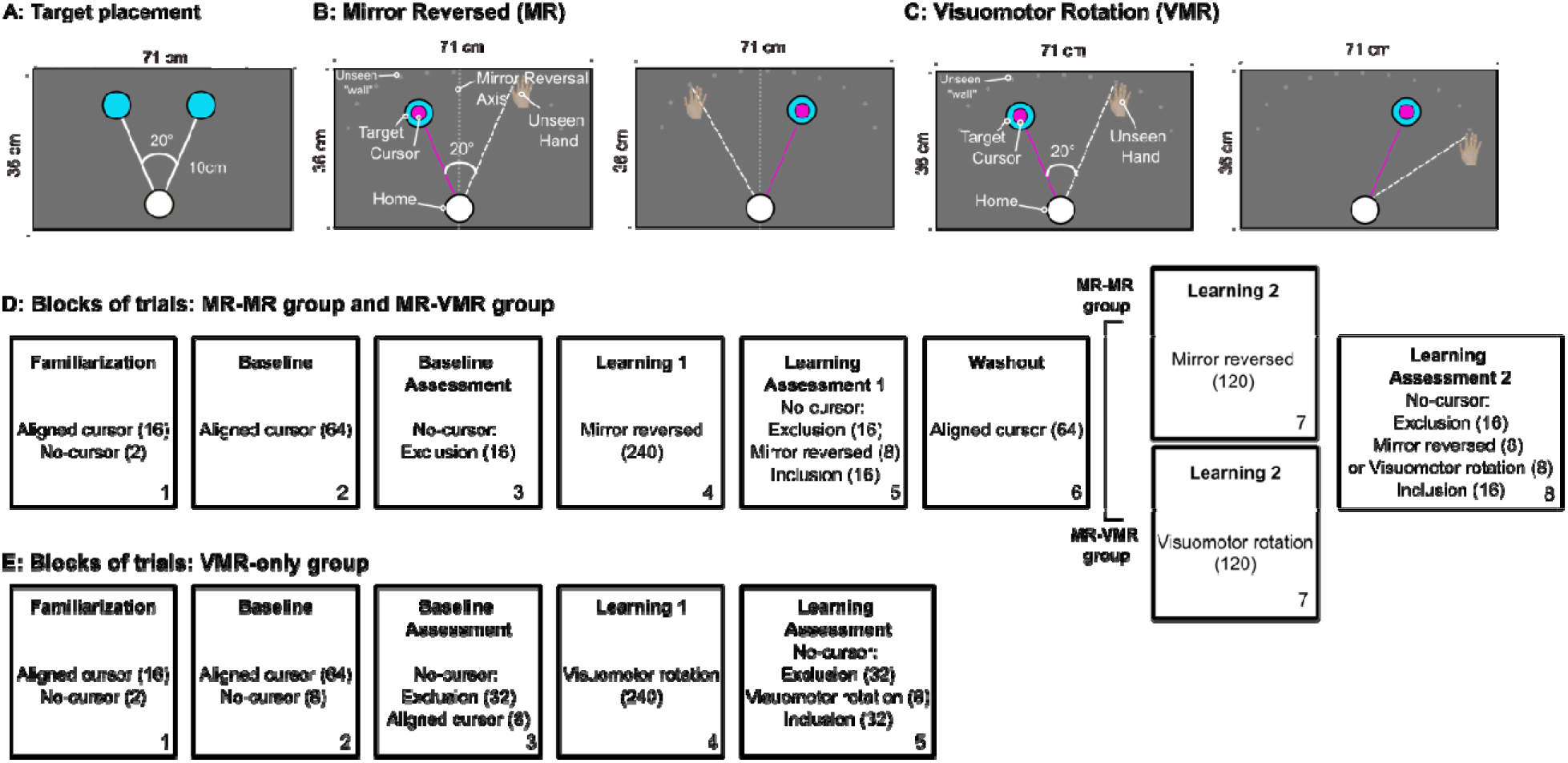
Target placement, visuomotor distortions and trial breakdown across blocks. **A:** The right target was positioned 10° to the right of the body midline (y-axis) and the left target was positioned 10° to the left of the body midline. **Training trials. B:** For the mirror reversed distortion, cursor motion was mirrored across the body midline (y-axis; light grey dashed line) relative to the index finger motion. When the left (right) target was shown, the index finger had to move 20° to the right (left) of the target for the cursor to land on the target. The dotted line depicts the mirroring axis, which was not visible to participants. **C:** For the visuomotor rotation distortion, cursor motion was rotated 20° clockwise (CW) or counterclockwise (CCW) relative to the index finger motion. For the CCW rotation shown in C, participants had to reach 20° to the right of the target for the cursor to land on both right and left targets. **D:** Breakdown of blocks of trials and the number of trials completed (in parentheses) within each block for the MR-MR and MR-VMR groups and **E:** for the VMR-only group.

Each trial began with participants positioning their index finger at the home position for 500 ms. This was followed by the appearance of both the home position and the cursor, followed by a variable delay of 300 to 700 ms before the target appeared. Participants performed rapid “slicing” movements through the target, aiming to intersect it at approximately peak velocity, consistent with ballistic shooting movements as described by Maksimovic et al. (2020). Movements were terminated by a soft mechanical wall (spring constant: 150 N/m; *unseen wall;* Figure 1B and C semicircular dotted line), placed 2 cm beyond the target array to encourage natural movement dynamics and promote rapid execution. Cursor feedback of the index finger’s position was available until the finger crossed the target array, after which the cursor disappeared. Movement onset was defined online as the time when the center of the cursor moved 0.5 cm from the center of the home position and the cursor’s velocity rose above 0.03 m/s and remained above that threshold for at least 9 ms. Movement end was recorded when the participant’s finger crossed the target array, and movement time (MT) was calculated as the time interval between movement onset and movement end.

Participants received MT feedback at the end of each trial via a change in target color: green indicated the goal MT (< 600 ms) had been achieved, whereas red indicated MT exceeded 600 ms. MT feedback was presented while the hand remained against the soft wall for 500 ms. The Kinarm then returned the finger to the home position along a straight trajectory over 1000 ms. This return movement occurred without visual feedback (see Figure 2).

**Figure 2.**
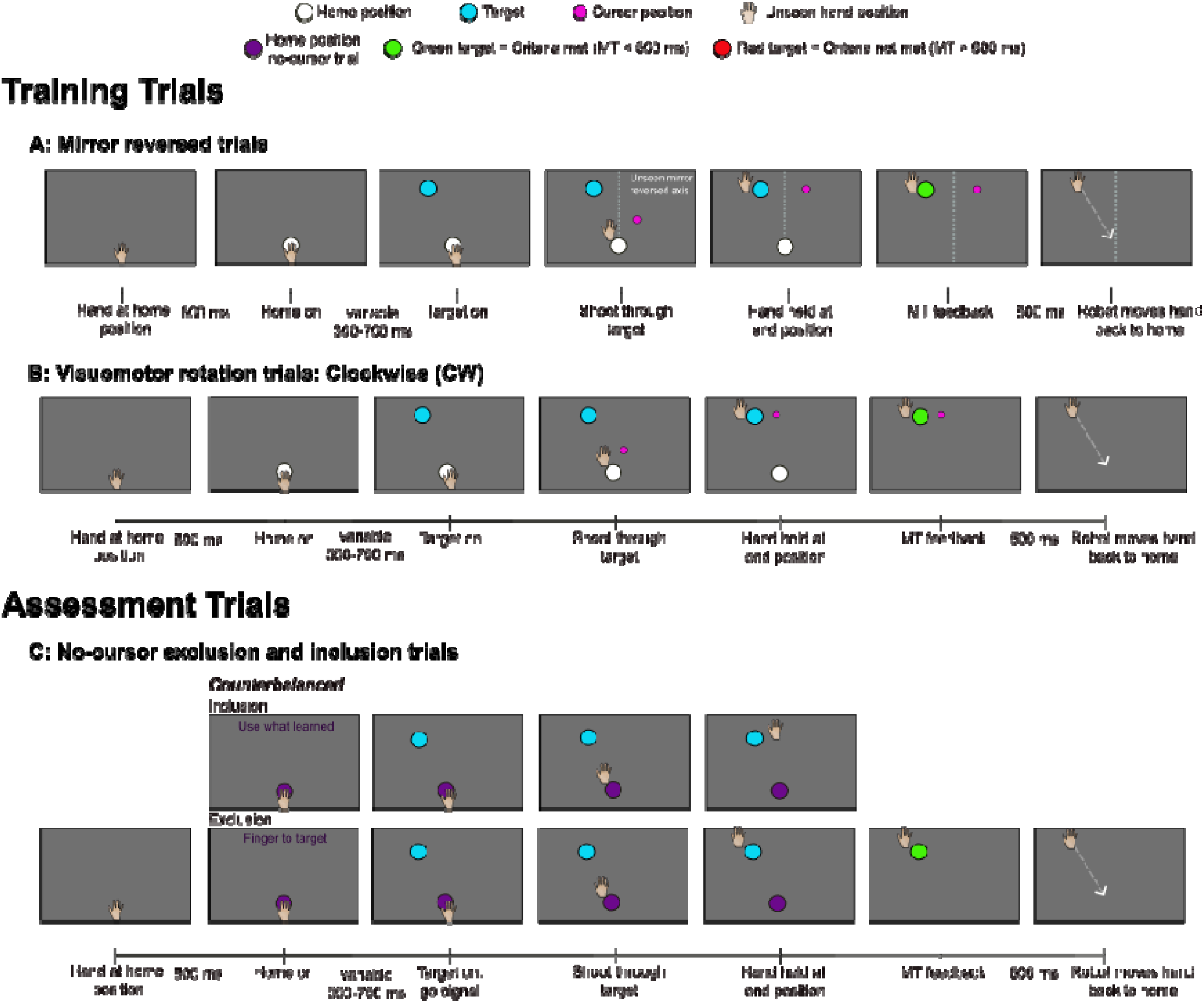
Timeline for trials completed. The hand and the index finger were hidden from participants’ view. **Training Trials. A:** Mirror reversed trials: The cursor’s trajectory was mirrored across the y-axis relative to the trajectory of the index finger and was visible throughout the reach until the finger crossed the target array. **B:** Visuomotor rotation trials: The cursor’s trajectory was rotated 20° CW or 20° CCW relative to the trajectory of the index finger and was visible throughout the reach until the finger crossed the target array. **Assessment Trials. C:** No-cursor PDP assessment trials: Reaches without cursor feedback. Inclusion (top row) and Exclusion (bottom row) trials.

### Types of Trials

#### Training trials

Participants performed slicing movements under one of three cursor feedback conditions. In aligned cursor trials, the position of a white cursor (0.5 cm in diameter) on the screen corresponded to the index finger’s position, providing veridical visual feedback. In mirror reversed feedback trials, the cursor mirrored the participant’s finger position across the y-axis, introducing a visuomotor distortion (Figure 1B; MR group). In trials with a visuomotor rotation, the cursor trajectory was rotated either 20° CW or CCW relative to actual hand motion (Figure 1C; VMR-CW and VMR-CCW groups). In learning block 1, the mirror reversed distortion was introduced (Figure 1D, block 4). Learning block 2 consisted of mirror reversed or visuomotor rotation distortion trials depending on the group (Figure 1D, block 7). The VMR-only group completed one block of learning trials with the visuomotor rotation distortion (Figure 1E, block 4). Participants were not instructed about the nature of the distortions.

#### Washout trials

Washout trials (Figure 1D, block 6) were completed after learning assessment block 1 and before learning block 2. The washout block was identical to the baseline block, such that participants completed slicing movements with aligned cursor feedback. The purpose of the washout block was to eliminate residual effects of reaching with the mirror reversed distortion, ensuring that participants returned to baseline levels of performance before beginning the second learning block.

#### Assessment trials - PDP: Exclusion and Inclusion trials

Participants completed these no-cursor trials under two instruction sets: exclusion and inclusion. Exclusion trials were used to establish implicit learning according to the Process Dissociation Procedure (PDP; adapted from Werner et al., 2015). For these trials, the words “Finger to Target” appeared above the target and participants were instructed:

*“You are now going to reach when you cannot see your* index finger*, as there will be no-cursor on the screen. Do not use anything you may have learned for these trials to get the cursor to the target. Instead, aim so that your* index finger *goes straight through the target as you did during baseline reaches.”*

Inclusion trials were used to establish explicit learning in accordance with the PDP method. For these no-cursor trials, the words “Use what learned” appeared above the target and participants were instructed:

*“You are now going to reach when you cannot see your index finger, as there will be no-cursor on the screen. For these trials, use anything you have learned during training to get the cursor to the target. In other words, aim so that the cursor would have gone straight to the target, as in the training trials you just completed.”*

The absence of the cursor in these trials was cued with a change in colour of the hom position, such that on these trials the home position was purple.

#### Data Analyses

All reaching trials were analyzed using custom-written MATLAB scripts (R2024a; The MathWorks, Inc.). The primary dependent variable was hand angle, defined as the angular difference between a reference vector (home position to target) and the reach endpoint vector (home position to movement end, defined as the finger position when the index finger crossed the target array). Hand angles at the position at which participants’ hands crossed the target array are reported. Given the ballistic nature of the shooting task, these hand angles correspond to hand angles at peak velocity. Reaction time (RT) was calculated as the time from target onset to movement onset (defined as when the cursor moved 0.5 cm away from the home position and velocity exceeded 0.03 m/s for at least 9 ms). Movement time (MT) was computed as the time from movement onset to movement end.

#### Outlier Detection

Five variables were examined to establish outlier trials: Start X, Start Y (initial x and y coordinates), hand angle, RT and MT. A trial was discarded if Start X, Start Y, or hand angle exceeded 3 standard deviations from a participant’s mean value for the same trial type within a given block. For example, each aligned cursor trial was compared to the mean of aligned cursor trials within the same block, while no-cursor trials were assessed relative to their block-specific mean. With respect to RT and MT, any trials with RT or MT less than 100 ms were excluded, as well as trials in which MT was longer than 600 ms. Overall, 17,520 trials were collected, of which 1,286 trials (7.3%) were excluded from analysis. Importantly, including or excluding these trials did not alter the overall statistical outcomes or the main effects reported below.

### Learning

#### Learning to reach with a mirror reversed distortion

The baseline and learning blocks were divided into early and late trials (first versus last 24 trials). To confirm that learning to reach with the mirror reversed distortion during early trials did not differ between groups, we compared hand angles in learning block 1 (Figure 1D, block 4) across the MR-MR and MR-VMR groups. Hand angles in learning block 1 were normalized by subtracting baseline hand angles for the same target and time (early versus late trials), yielding normalized learning values for group comparisons. The absolute values of normalized hand angles were then compared across groups in a 2 group (MR-MR, MR-VMR) x 2 time (early, late) mixed analysis of variance (ANOVA) with repeated measures (RM) on the last factor. A similar analysis was completed for hand angle variability, RT and MT, though these data were not normalized to baseline data. Data related to learning to reach with the mirror reversed distortion for the MR-VMR group is presented in the Supplementary File.

#### Washout

We analyzed hand angles during the washout trials in a 2 group (MR–MR, MR–VMR) × 3 time (late baseline, early washout, late washout) mixed ANOVA with RM on the last factor to detect evidence of implicit learning.

#### Re-learning to reach with an mirror reversed distortion

To examine savings in the MR–MR group, we compared performance during the first and second block of trials of reaching with the mirror reversed distortion. Specifically, a 2 block (learning block 1, learning block 2) x 2 time (early, late trials) RM ANOVA was conducted on normalized hand angles. The same analyses were conducted for hand angle variability, RT and MT within the same blocks.

#### Learning to reach with a visuomotor distortion

To examine whether learning to reach with a mirror reversed distortion influenced learning to reach with a visuomotor rotation distortion, we compared normalized hand angles for reaches with the visuomotor rotation distortion between the MR–VMR group and the VMR-only group. Specifically, a 2 group (MR-VMR group, VMR-only group) x 2 time (early, late) mixed ANOVA was conducted with RM on the last factor. The same analyses were conducted for hand angle variability, RT and MT. To further examine potential group differences early in training with the visuomotor rotation distortion that may have been obscured when averaging across the first 24 trials, the early 24 trials were subdivided into three consecutive bins of 8 trials (Bin 1: trials 1–8; Bin 2: trials 9–16; Bin 3: trials 17–24). Means of these 8 trials were then normalized by the mean of the corresponding 8 trial bin in the baseline block and analyzed in a 2 group (MR-VMR, VMR-only) × 3 bin (Bin 1, Bin 2, Bin 3) mixed ANOVA with RM on the last factor.

#### Implicit and Explicit learning

Implicit and explicit learning were assessed via the PDP exclusion and inclusion trials following learning block 1 (Figure 1D & E, block 4) and learning block 2 (Figure 1D, block 8). All values were normalized relative to the corresponding PDP trials in the baseline block, such that positive values indicated a deviation in the expected direction of learning (i.e., + 20°) for both implicit and explicit learning.

The implicit index (I_Ind_) was calculated for each target within the baseline and learning assessment blocks in accordance with the following formula:

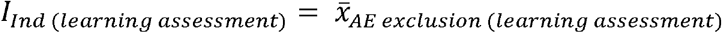

This index was then used to calculate implicit learning according to the following formula:

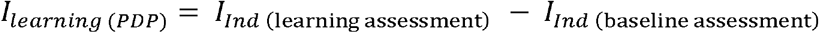

The explicit index (E_Ind_) for awareness of changes in reaches was established according to the following formulas:

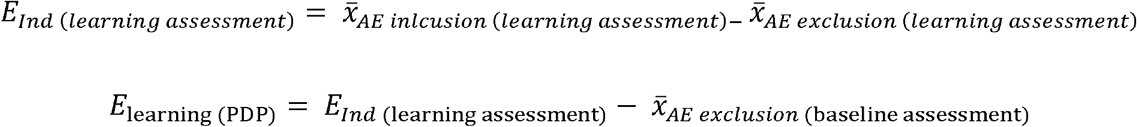

To determine whether implicit and explicit learning were reliably expressed, each group’s implicit and explicit learning were compared to zero using one-sample t-tests. Then, to assess the engagement of implicit and explicit processes at different stages of learning (i.e., learning assessment block 1 and learning assessment block 2) within and between groups, three different analyses were completed. First, implicit and explicit learning following learning block 1 were compared between groups (MR–MR, MR–VMR) using independent *t*-tests. Second, within the MR–MR group, implicit and explicit learning in learning block 1 and learning block 2 were compared using paired-samples *t*-tests. Third, to determine whether learning to reach with a mirror reversed distortion influenced the processes engaged during subsequent reaches with the visuomotor rotation distortion, implicit and explicit learning following learning block 2 were compared between the MR–VMR and VMR-only groups using independent *t*-tests.

All statistical analyses were conducted using JASP, RStudio, and Microsoft Excel. In cases where Levene’s test indicated a violation of homogeneity of variance, a Welch correction was applied. For ANOVA analyses, sphericity was assessed using Mauchly’s test and Greenhouse-Geisser corrections were applied when appropriate. The alpha level for significance was set at *p* < 0.05, and Bonferroni corrections were used for all post hoc tests. Effect sizes are reported as partial eta squared (η*_p_²*) or Cohen’s d (*d*). For clarity, results focus on the highest-order significant interactions and lower-order effects are reported only when relevant.

## Results

### Learning

#### Learning to reach with a mirror reversed distortion

With respect to hand angles when the MR distortion was introduced in learning block 1, ANOVA revealed a significant main effect of time (*F*(1,23) = 75.318, *p* < 0.001, η*_p_²* = 0.766). Post hoc analyses indicated that early hand angles were significantly smaller than late hand angles (both *p* < 0.001). There was no significant main effect of group (*F*(1,23) = 0.165, *p* = 0.688, η*p²* = 0.007), nor a significant group × time interaction (*F*(1,23) = 0.031, *p* = 0.863, η*p²* = 0.001), indicating comparable learning across the MR-MR and MR-VMR groups (Figure 3). For hand angle variability, RT and MT, no significant main effects or interactions were observed (all *p* > 0.05; Figure 4C, D; Figure S1C, D).

**Figure 3.**
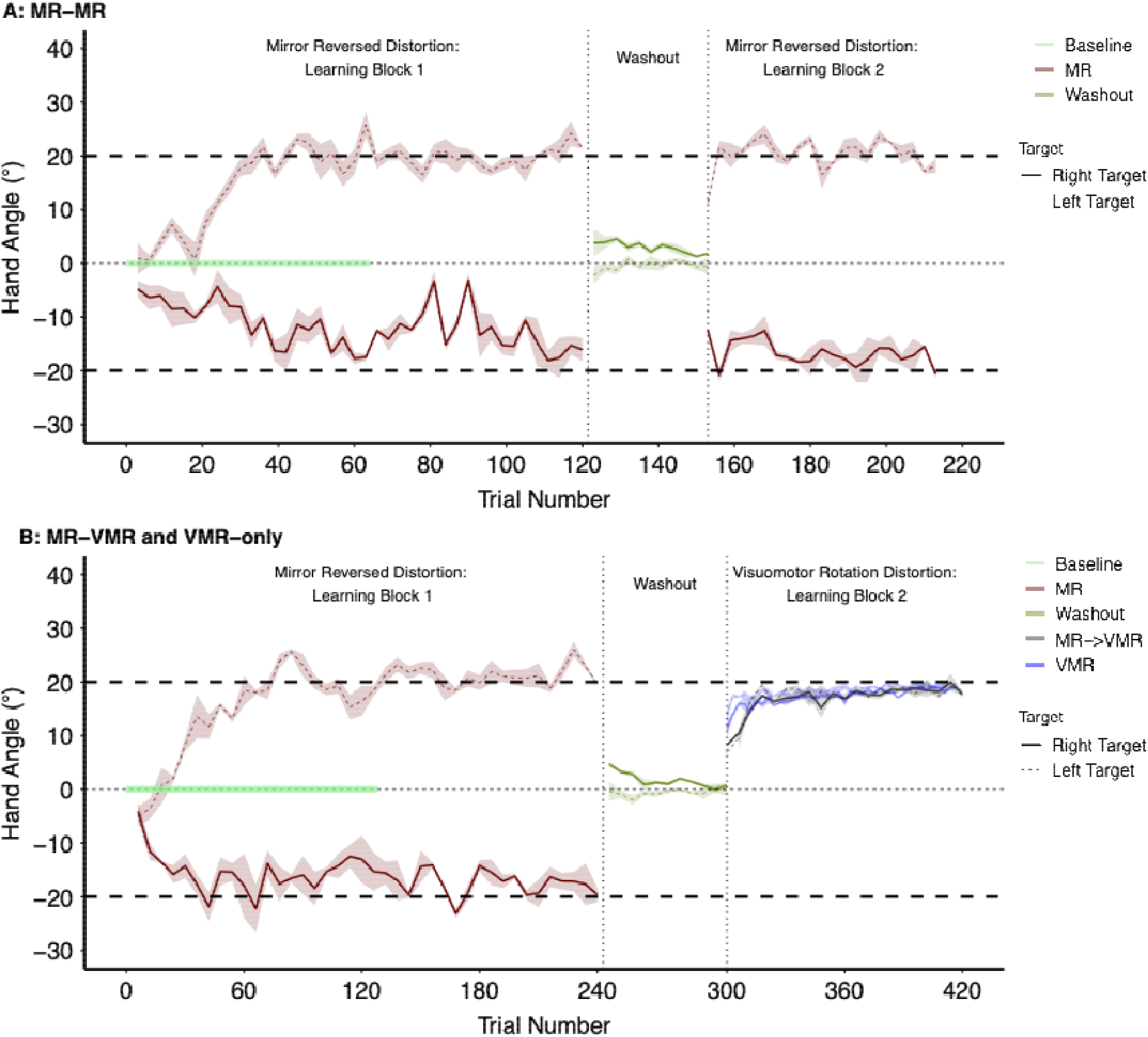
Mean hand angles for **A:** the MR-MR group and **B:** the MR-VMR group across all trials. Data from the VMR-only group are shown as blue traces during reaches with the visuomotor rotation distortion in panel B. For both panels, baseline and washout data are depicted in different shades of green. Solid lines represent reaches to the right target (+20°), whereas dashed lines represent reaches to the left target (−20°). During learning blocks with the visuomotor rotation distortion, reaches to both targets converge towards a similar hand angle (+20). Dashed horizontal lines indicate complete learning. Thin dashed vertical lines mark the boundaries between experimental blocks.

**Figure 4.**
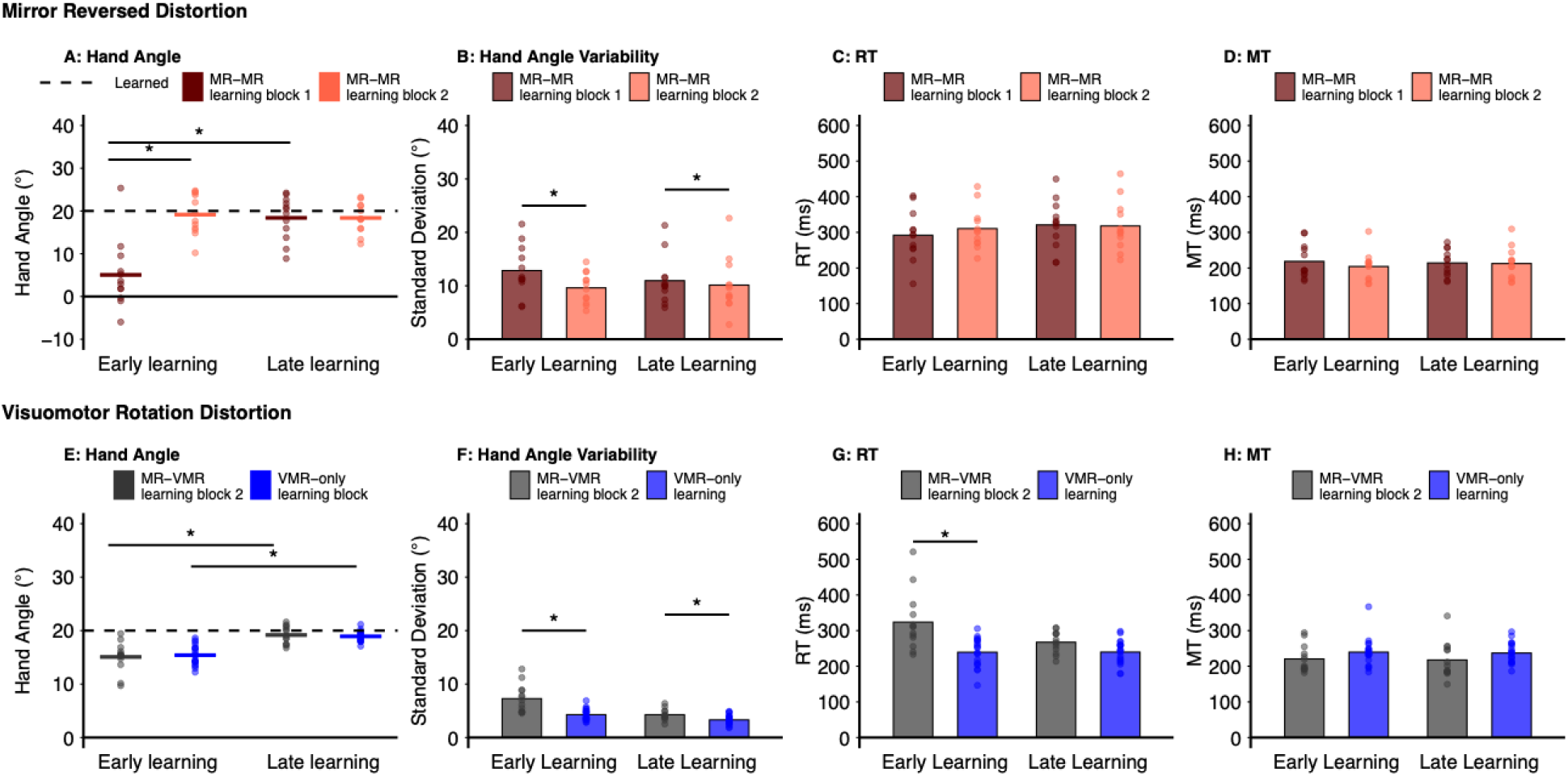
Hand angles, hand angle variability, reaction time (RT) and movement time (MT) during reaches with the mirror reversed and visuomotor rotation distortions. **A–D:** Performance during early and late trials when reaching with the mirror reversed distortion for the MR–MR group in learning block 1 (dark red) and learning block 2 (light red). **A:** Hand angle. **B:** Hand angle variability (standard deviation). **C:** RT. **D:** MT. **E–H:** Performance during early and late trials when reaching with the visuomotor rotation distortion for the MR–VMR (grey) and VMR-only (blue) groups. **E:** Hand angle. **F:** Hand angle variability. **G:** RT. **H:** MT. Dots represent individual participant data, while bars and lines indicate group means. Asterisks indicate statistically significant differences within or between blocks for the MR-MR group (top row) or between groups (bottom row; *p* < 0.05).

#### Washout

No significant main effects or interactions were observed (all *p* > 0.05), indicating that hand angles remained comparable across the washout trials and between groups. Hand angles were comparable to baseline for both the MR-MR and MR-VMR groups during early and late washout trials (Figure 3A, B; MR-MR: late baseline M = 0.4° ± 0.9°, early washout: M = 1.4° ± 1.7°; late washout M = 0.8° ± 0.8°; MR-VMR group: late baseline M = 0.002° ± 1.0°, early washout: M = 0.9° ± 2.4°; late washout M = 0.3° ± 1.1°), suggesting that implicit proccesses were not engaged during the washout block.

#### Re-learning to reach with a mirror reversed distortion

To assess for savings in the MR–MR group, early and late hand angles in learning block 1 (Figure 1, block 4) were compared with those in learning block 2 (Figure 1, block 7). ANOVA revealed significant main effects of block (*F*(1,11) = 30.863, *p* < 0.001, η*_p_²* = 0.737), time (*F*(1,11) = 46.829, *p* < 0.001, η*_p_²* = 0.810), and a significant block x time interaction (*F*(1,11) = 34.985, *p* < 0.001, η*_p_²* = 0.761). Post hoc analyses indicated that hand angles were larger (i.e., closer to complete learning) in the early trials in learning block 2 compared to the early trials in learning block 1, indicating savings. Early hand angles in learning block 2 did not differ from late hand angles in the same block or late hand angles in learning block 1 (*p* > 0.05; Figure 4A), indicating no further changes across learning block 2.

With respect to hand angle variability in the MR-MR group, ANOVA revealed a significant main effect of block (*F*(1,11) = 10.915, *p* = 0.007, η*_p_²* = 0.498). Post hoc comparisons revealed that hand angle variability was greater in learning block 1 compared to learning block 2 (*p* = 0.007; Figure 4B). RT and MT did not differ between learning block 1 and learning block 2 (all *p* > 0.05; Figure 4C, D).

#### Learning to reach with the visuomotor rotation distortion

We compared learning to reach with the visuomotor rotation distortion in the MR-VMR group to the VMR-only group. When 24 trials were included within the early and late trial means, ANOVA revealed a significant main effect of time (*F*(1, 31) = 71.105, *p* < 0.001, η*_p_²* = 0.696). Hand angles in the early trials were less than hand angles in the late trials for each group (Figure 4E). No significant main effects or interactions involving group were observed (all *p* > 0.05). However, when the analysis was repeated with the factor of bin included, such that only 8 trials were included in the mean for early trial hand angles for each bin, ANOVA revealed a significant main effect of bin (*F*(1.550, 48.047) = 70.312, *p* < 0.001, η*_p_²* = 0.694) and a significant group × bin interaction (*F*(1.550, 48.047) = 10.351, *p* < 0.001, η*_p_²* = 0.250). Post hoc analyses indicated that hand angles during the first bin (trials 1–8) tended to be smaller in the MR–VMR group (M = 8.8° ± 5.4°) compared to the VMR-only group (M = 12.6° ± 3.3°), although this difference did not reach statistical significance (*p* = 0.051). Hand angle in the second and third bin of 8 trials did not differ between groups (all *p* > 0.05; Figure 3B, see Visuomotor Rotation Distortion: Learning Block 2).

With respect to hand angle variability, ANOVA revealed a main effect of group (*F*(1, 31) = 4.472, *p* = 0.047, η*_p_²* = 0.420), time (*F*(1, 31) = 41.976, *p* < 0.001, η*_p_²* = 0.575), and a significant group x time interaction (*F*(1, 31) = 12.933, *p* = 0.001, η*_p_²* = 0.294). Post hoc analyses indicated that hand angle variability was greater when learning to reach with the visuomotor rotation distortion for the MR–VMR group compared to the VMR-only group for both early and late trials (all *p* < 0.05; Figure 4F).

RT also differed between the MR-VMR and VMR-only groups, with ANOVA revealing a significant main effect of group (*F*(1, 31) = 14.963, *p* < 0.001, η*_p_²* = 0.326), and time (*F*(1, 31) = 9.719, *p* = 0.004, η*_p_²* = 0.239), and a significant group x time interaction (*F*(1, 31) = 9.986, *p* = 0.004, η*_p_²* = 0.244). Post hoc analyses indicated that the MR-VMR group reached with longer RTs than the VMR-only group during early trials (*p* = 0.002; Figure 4G). In contrast, RTs did not differ between groups during late trials (*p* = 0.124; MR-VMR: 267 ms; VMR-only: 240 ms). MT did not differ between groups or between early and late learning trials (all *p* > 0.05; Figure 4H).

#### Implicit learning

Implicit learning is displayed in Figure 5 following learning to reach with the mirror reversed distortion in learning block 1. Implicit learning did not differ between the MR–MR and MR–VMR groups (*t*(23) = −1.224, *p* = 0.233, *d* = −0.490). Within the MR–MR group, implicit learning did not change from learning block 1 to learning block 2 (*t*(11) = −1.101, *p* = 0.294), indicating comparable levels of implicit learning across reaches with the mirror reversed distortion (i.e., in learning blocks 1 and 2). Although implicit estimates following reaches with the mirror reversed distortion differed significantly from zero (all *p* < 0.05), the direction of these estimates was opposite to that expected of learning.

**Figure 5.**
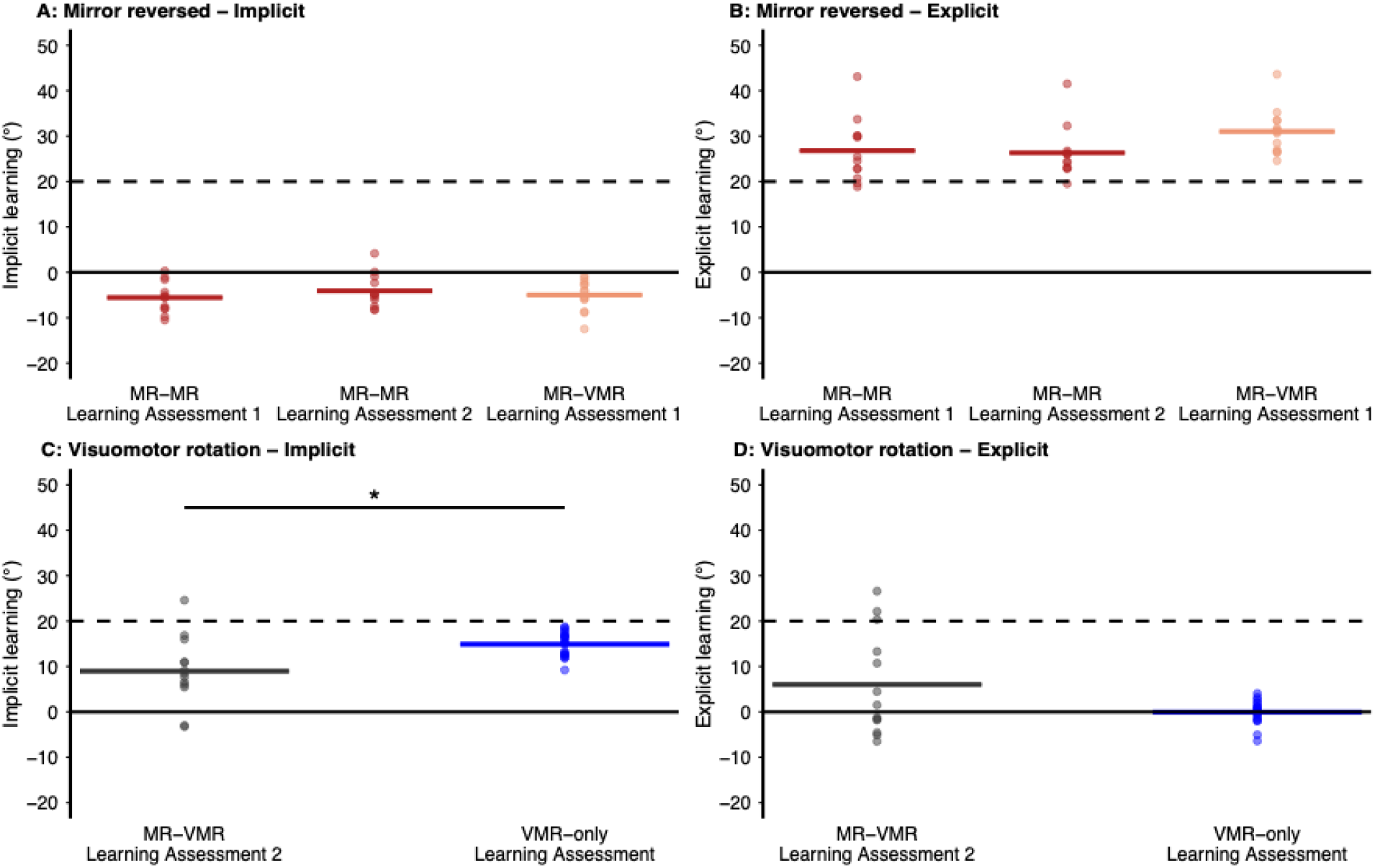
Implicit and explicit learning following reaches with the mirror reversed and visuomotor rotation distortions. **A:** Implicit learning and **B:** Explicit learning following reaches with the mirror reversed distortion for the MR-MR group (dark red) in learning block 1 and learning block 2, and for the MR-VMR group (light red) in learning block 1. **C:** Implicit learning and **D:** Explicit learning following reaches with the visuomotor rotation distortion for the MR-VMR group (grey) in learning block 2 and for the VMR-only group (blue). Circles represent individual participant data; bold lines indicate group means. Dashed lines indicate the expected direction of learning (+20°). Asterisks indicate significant differences between groups (*p* < 0.05).

Implicit learning following reaches with the visuomotor rotation distortion differed between the MR-VMR and VMR-only groups (*t*(31) = −4.532, *p* < 0.001, *d* = 0.477), with smaller implicit learning observed for the MR-VR group (MR–VMR learning block 2: M = 8.8° ± 5.1°) compared to the VMR-only group (M = 14.9° ± 2.6°). For both groups, implicit learning was significantly greater than zero (all *p* < 0.05), in the expected direction of learning.

#### Explicit learning

Explicit learning is shown in Figure 5. Following initial reaches with the mirror reversed distortion (learning block 1), explicit learning did not differ between the MR–MR and MR–VMR groups (*t*(23) = −1.762, *p* = 0.091, *d* = −0.705). Within the MR–MR group, explicit learning did not differ between learning block 1 and learning block 2 (*t*(11) = 0.439, *p* = 0.669), indicating similar explicit learning across reaches with the mirror reversed distortion (i.e., in learning blocks 1 and 2). Explicit learning was significantly greater than zero following all reaches with the mirror reversed distortion (all *p* < .05; MR–MR learning block 1: M = 26.8° ± 7.0°; MR–MR learning block 2: M = 25.9° ± 6.5°; MR–VMR learning block 1: M = 31.0° ± 5.0°), and in the expected direction of learning.

Explicit learning following reaches with the visuomotor rotation distortion did not differ between the MR-VMR and VMR-only groups, *t*(12.8) = 1.901, *p* = 0.080, *d* = 0.739. Although the between-group comparison did not reach statistical significance, the effect size indicated a moderate difference, with greater explicit learning in the MR–VMR group (M = 6.0° ± 11.6°) than in the VMR-only group (M = 0.07° ± 2.6°). Moreover, explicit learning within the MR–VMR group was larger than zero in the expected direction of learning (*p* < 0.001), whereas explicit learning in the VMR-only group did not differ from zero (*p* = 0.911).

## Discussion

The present study examined whether the processes engaged during initial learning influence subsequent motor learning. Specifically, we asked whether learning to reach with a small mirror reversed distortion, which is primarily supported by explicit processes, supports savings when the same distortion is re-introduced. We further asked if learning to reach with a mirror reversed distortion influences learning to reach with a small visuomotor rotation distortion, which has been shown to be driven by implicit processes (Neville & Cressman, 2018; Werner et al., 2015). We observed clear evidence of savings in our MR–MR group, with participants demonstrating reduced initial hand angles when re-learning to reach with the mirror reversed distortion (i.e., learning block 2), suggesting that participants were able to rapidly reinstate explicit strategies acquired during initial learning (i.e., learning block 1). There was a trend for learning to reach with the mirror reversed distortion to influence reaches when the visuomotor rotation distortion was first introduced, such that hand angles were smaller (i.e., demonstrated less complete compensation for the cursor rotation) when reaches with the visuomotor rotation distortion were preceded by mirror reversed reaches. As well, early learning trials with the visuomotor rotation distortion in the MR-VMR group demonstrated greater reach variability and longer reaction times (RTs) compared to the VMR-only group. Finally, prior reaches with the mirror reversed distortion significantly altered the underlying learning processes engaged when learning to reach with the visuomotor rotation distortion such that the MR–VMR group demonstrated engagement of explicit processes and the engagement of implicit processes was reduced relative to the VMR-only group.

### Savings when reaching with a mirror reversed distortion reflects retrieval of explicit processes

The present findings provide evidence of savings when re-learning to reach with a small mirror reversed distortion. Although savings is typically characterized as a faster rate of re-learning, we did not observe steeper learning curves with respect to error reduction in learning block 2 compared to block 1 for our MR-MR group. Rather, participants began re-learning at a level comparable to their late performance in learning block 1. Consequently, savings was expressed as reduced initial errors upon re-exposure to the distortion rather than as an increased rate of improvement.

While previous research has demonstrated persistence of learning when reaching with a mirror reversed distortion across three days (Gastrock et al., 2024), and offline performance gains following sleep (Telgen et al., 2014), these studies did not directly assess savings upon re-learning to reach with the same mirror reversed distortion within a single testing session. Gastrock and colleagues (2024) showed that reach completion time (defined as the time elapsed from target onset to target acquisition, encompassing RT and movement time, MT) and movement path length were shorter at the start of a second testing session three days after a first session, indicating retention of learning to reach with the mirror reversed distortion across three days. Similarly, Telgen et al. (2014) demonstrated that early reaching performance with a mirror reversed distortion following a night of sleep was equal to or better than performance at the end of the preceding training session, suggesting offline gains in learning to reach with a mirror reversed distortion. The current results extend this prior work by demonstrating savings upon re-learning to reach with a mirror reversed distortion within a single testing session.

Prior work has demonstrated that learning to reach with a mirror reversed distortion is driven primarily by explicit processes, with little to no implicit contributions (Wang & Taylor, 2021; Wilterson & Taylor, 2021; Hadjiosif et al., 2021; Heirani Moghaddam, Manson, Cressman, submitted). Consistent with this, we observed no evidence of implicit processes contributing to savings when re-learning to reach with the mirror reversed distortion. Instead, explicit learning was evident following reaches with the mirror reversed distortion in both learning block 1 and learning block 2, and the magnitude of explicit learning did not differ between blocks.

The observation that explicit processes supported savings parallels prior demonstrations of savings during learning to reach with a visuomotor rotation distortion (Krakauer et al., 2019; Leow et al., 2016; Morehead et al., 2015), where savings has been proposed to arise due to rapid re-engagement of explicit reach strategies. For example, Morehead et al. 2015 (see also Werner et al., 2015) showed that savings in learning to reach with a visuomotor rotation distortion was observed when explicit processes were engaged during initial learning. In contrast, implicit processes do not appear to reliably support savings when reaching with a visuomotor rotation distortion (Bond & Taylor, 2015). Werner et al. (2015) reported evidence of savings following learning to reach with larger cursor rotations (e.g., 40° and 60°), that engage both implicit and explicit processes, but not following learning to reach with a 20° cursor rotation, which is supported by implicit processes (Neville & Cressman, 2018).

Taken together, the present findings extend observations from the visuomotor rotation literature by suggesting that savings following learning to reach with a small mirror reversed distortion also reflects retrieval of explicit processes in the absence of substantial implicit contributions. Savings observed in the MR-MR group was accompanied by reduced reach variability, without increases in RT or MT. This suggests efficient reinstatement of an explicit solution rather than additional planning and execution demands. Overall, these results demonstrate that learning to reach with a small mirror reversed distortion exhibits savings within a single testing session, suggesting that once established, the explicit strategy used when learning to reach with a mirror reversed distortion can be efficiently reinstated.

### Learning to reach with a mirror reversed distortion interferes with learning to reach with a visuomotor rotation distortion

Interference in learning to reach with a visuomotor rotation distortion has commonly been observed when participants reach with opposing or competing visuomotor rotation distortions (e.g., reaching with a CW cursor rotation following learning to reach with a CCW cursor rotation), where learning of one visuomotor mapping impairs acquisition or retention of a subsequent visuomotor mapping (Brashers-Krug et al., 1996; Krakauer et al., 2005). In these paradigms, interference is typically expressed behaviourally as increased initial errors, slower learning rates, or reduced retention (Caithness et al., 2004; Krakauer et al., 2005). Prior work suggests that interference when learning to reach with a visuomotor rotation distortion primarily reflects disruption or competition at the level of explicit processes, particularly when previously established strategies must be suppressed or replaced (Haith et al., 2015; McDougle & Taylor, 2019).

We found that learning to reach with a mirror reversed distortion did not alter the extent of learning achieved when reaching with a visuomotor rotation distortion in late learning trials, as reflected by comparable late hand angles between the MR–VMR and VMR-only groups. Gastrock and colleagues (2024) have reported similar findings, such that prior reaches with a mirror reversed distortion did not facilitate or impair learning to reach with a visuomotor rotation distortion. That said, analysis of early reaches with the visuomotor rotation distortion in our study revealed a trend towards smaller hand angles in the first 8 trials when the visuomotor rotation distortion was first introduced for the MR-VMR group compared to the VMR-only group. Further, we found that reaches with the visuomotor rotation distortion following reaches with the mirror reversed distortion were characterized by greater reach variability and longer RT early in training. These findings suggest that prior reaches with the mirror reversed distortion interfered with learning to reach with the visuomotor rotation distortion.

One explanation for this interference is that the explicit solution established during learning to reach with the mirror reversed distortion continued to be engaged when the visuomotor rotation distortion was introduced. To test this possibility, we examined whether learning to reach with the mirror reversed distortion altered the underlying processes engaged during learning to reach with the visuomotor rotation distortion. We found that learning to reach with a small mirror reversed distortion altered the processes engaged during reaches with the visuomotor rotation distortion. Specifically, implicit contributions were reduced in the MR–VMR group relative to the VMR-only group. In addition, explicit processes were observed following reaches with the visuomotor rotation distortion in the MR–VMR group, whereas explicit learning did not differ from zero in the VMR-only group. In other words, the MR-VMR group demonstrated greater explicit learning following reaches with the mirror reversed distortion compared to reaches with the visuomotor rotation distortion. Implicit learning revealed the opposite trend, such that implicit learning was greater following reaches with the visuomotor rotation distortion compared to the mirror reversed distortion.

Interestingly, in the MR–VMR group, implicit learning following reaches with the MR distortion differed from zero, but was in the direction opposite to that of expected learning. These results are in agreement with Hadjosif et al. (2021), and suggest that the observed changes in reaches do not reflect meaningful implicit learning. Instead, this pattern of results suggests that while implicit processes continue to be updated in response to sensory prediction errors during reaches with a mirror reversed distortion, these updates occur in a direction that does not support successful compensation for the distortion. As such, these effects may reflect inappropriate or nonspecific movement updating rather than functionally relevant implicit learning.

Together, these findings indicate that learning to reach with a mirror reversed distortion interfered with the underlying processes engaged when learning to reach with a visuomotor rotation distortion. This suggests that initial engagement of explicit processes may establish a durable control policy, biasing subsequent learning towards explicit processes even when the new visuomotor distortion would typically be supported by implicit processes. Thus, prior learning can shape the allocation of learning processes in future contexts. In this sense, prior experience of learning to reach with a mirror reversed distortion appears to restructure how the system coordinates implicit and explicit contributions when a new visuomotor mapping is encountered, accommodating prior learning by continuing to engage explicit processes.

### Methodological considerations and limitations

The present findings support the suggestion that the processes engaged during initial learning influence subsequent motor learning. That said, several methodological considerations should be acknowledged when interpreting our findings and designing future research. Methodological considerations include the number of training and PDP trials completed by each group, as well as the number of targets that participants reached to. The MR–VMR group completed additional reach training trials prior to reaching with the visuomotor rotation distortion, potentially increasing their level of fatigue compared to the VR-only group. That said, the observed results argue against fatigue influencing performance. For example, in the MR–VMR group both reach variability and RT decreased from early to late learning trials in learning block 2, indicating improved performance over time rather than a progressive decline. This is inconsistent with the suggestion of fatigue influencing performance, as fatigue-related effects would be expected to accumulate with continued task exposure and lead to decreased performance (Behrens et al., 2023; Dallaway et al., 2022). In addition, RT for the VMR-only group at the end of the baseline block did not differ from RT in early learning trials in the learning block, suggesting that the transition from reaches with an aligned cursor to visuomotor rotation distortion did not lead to elevated RT. Together, these findings argue against fatigue driving the observed group differences and instead support the interpretation that learning to reach with a mirror reversed distortion altered the processes engaged during reaches with a visuomotor rotation distortion.

A second methodological consideration is that participants in the MR–VMR group completed the PDP trials twice, whereas participants in the VMR-only group completed the PDP trials only once following reaches with the visuomotor rotation. Repeated exposure to PDP instructions may increase awareness of task demands or encourage the use of explicit strategies. However, differences in reaction time and reach variability were evident during early learning trials with the visuomotor rotation, before the second PDP assessment occurred, suggesting that repeated completion of the PDP trials is unlikely to fully explain group differences.

A limitation of the present study is the use of only two target locations. While this design was intentionally selected to maintain a consistent distortion magnitude across the mirror reversed and visuomotor rotation distortions, it may have encouraged the development of target-specific solutions rather than learning of a visuomotor mapping across a broader workspace. Although related work from our laboratory using a similar mirror reversed paradigm has demonstrated generalization to novel target locations, the present findings should be interpreted within the context of the target set employed. Future work should determine whether the patterns of savings and interference observed here generalize to paradigms involving a larger number of target locations. A final limitation of the present study is the modest sample size. While sensitivity analyses indicated that the design was adequately powered to detect medium-to-large within-subject effects and large between-group effects, smaller effects may have gone undetected.

## Conclusion

Overall, our findings demonstrate that learning to reach with a small mirror reversed distortion establishes a durable, explicitly mediated reaching strategy that can be rapidly reinstated when the distortion is re-introduced, even after washout trials. This reaching strategy not only supports savings during re-learning to reach with the mirror reversed distortion but also influences how learning to reach with a visuomotor rotation distortion unfolds, such that explicit processes are engaged.

## Ethics approval

This study was approved by the Queen’s University Health Sciences and Affiliated Teaching Hospitals Research Ethics Board (HSREB).

## Consent to participate

Informed consent was obtained from all individual participants included in the study.

## References

Avraham, G., Morehead, J. R., Kim, H. E., & Ivry, R. B. (2021). Reexposure to a sensorimotor perturbation produces opposite effects on explicit and implicit learning processes. PLoS Biology, 19(3), e3001147. 10.1371/journal.pbio.3001147

Baraduc, P., & Wolpert, D. M. (2002). Adaptation to a visuomotor shift depends on the starting posture. Journal of Neurophysiology, 88(2), 973–981. 10.1152/jn.2002.88.2.973

Bastian, A. J. (2008). Understanding sensorimotor adaptation and learning for rehabilitation. Current Opinion in Neurology, 21(6), 628–633. 10.1097/WCO.0b013e328315a293

Behrens, M., Gube, M., Chaabene, H., Prieske, O., Zenon, A., Broscheid, K.-C., Schega, L., Husmann, F., & Weippert, M. (2023). Fatigue and Human Performance: An Updated Framework. Sports Medicine (Auckland, N.z.), 53(1), 7–31. 10.1007/s40279-022-01748-2

Benson, B. L., Anguera, J. A., & Seidler, R. D. (2011). A spatial explicit strategy reduces error but interferes with sensorimotor adaptation. Journal of Neurophysiology, 105(6), 2843–2851. 10.1152/jn.00002.2011

Bock, O., Schneider, S., & Bloomberg, J. (2001). Conditions for interference versus facilitation during sequential sensorimotor adaptation. Experimental Brain Research, 138(3), 359–365. 10.1007/s002210100704

Bond, K. M., & Taylor, J. A. (2015). Flexible explicit but rigid implicit learning in a visuomotor adaptation task. Journal of Neurophysiology, 113(10), 3836–3849. 10.1152/jn.00009.2015

Bouchard, & Cressman, E. K. (2021). Intermanual transfer and retention of visuomotor adaptation to a large visuomotor distortion are driven by explicit processes. PLoS ONE, 16(1 January), 1–20. 10.1371/journal.pone.0245184

Brashers-Krug, T., Shadmehr, R., & Bizzi, E. (1996). Consolidation in human motor memory. Nature, 382(6588), 252–255. 10.1038/382252a0

Caithness, G., Osu, R., Bays, P., Chase, H., Klassen, J., Kawato, M., Wolpert, D. M., & Flanagan, J. R. (2004). Failure to consolidate the consolidation theory of learning for sensorimotor adaptation tasks. The Journal of Neuroscience: The Official Journal of the Society for Neuroscience, 24(40), 8662–8671. 10.1523/JNEUROSCI.2214-04.2004

Dallaway, N., Lucas, S., & Ring, C. (2022). Cognitive tasks elicit mental fatigue and impair subsequent physical task endurance: Effects of task duration and type. Psychophysiology, (e14126). 10.1111/psyp.14126

de Brouwer, A. J., Albaghdadi, M., Flanagan, J. R., & Gallivan, J. P. (2018). Using gaze behavior to parcellate the explicit and implicit contributions to visuomotor learning. Journal of Neurophysiology, 120(4), 1602–1615. 10.1152/jn.00113.2018

Desrochers, P. C., Brunfeldt, A. T., & Kagerer, F. A. (2020). Neurophysiological Correlates of Adaptation and Interference during Asymmetrical Bimanual Movements. Neuroscience, 432, 30–43. 10.1016/J.NEUROSCIENCE.2020.01.044

Gastrock, R. Q., ’T Hart, B. M., & Henriques, D. Y. P. (2024). Distinct learning, retention, and generalization patterns in de novo learning versus motor adaptation. Scientific Reports, 14(1), 8906. 10.1038/s41598-024-59445-1

Ghahramani, Z., Wolpert, D. M., & Jordan, M. I. (1996). Generalization to local remappings of the visuomotor coordinate transformation. The Journal of Neuroscience : The Official Journal of the Society for Neuroscience, 16(21), 7085–7096.

Haith, A. M., Huberdeau, D. M., & Krakauer, J. W. (2015). The influence of movement preparation time on the expression of visuomotor learning and savings. Journal of Neuroscience, 35(13), 5109–5117. 10.1523/JNEUROSCI.3869-14.2015

Heirani Moghaddam, S., Cressman, E. K., & Manson, G. A. (2026). Implicit processes do not contribute to learning to reach in small mirror reversed visuomotor environments. PLOS ONE, 21(6), e0333564. 10.1371/journal.pone.0333564

Izawa, J., Criscimagna-Hemminger, S. E., & Shadmehr, R. (2012). Cerebellar Contributions to Reach Adaptation and Learning Sensory Consequences of Action. Journal of Neuroscience, 32(12), 4230–4239. 10.1523/JNEUROSCI.6353-11.2012

Izawa, J., & Shadmehr, R. (2011). Learning from sensory and reward prediction errors during motor adaptation. PLoS Computational Biology, 7(3), 1–11. 10.1371/journal.pcbi.1002012

Jacoby, L. L. (1991). A process dissociation framework: Separating automatic from intentional uses of memory. Journal of Memory and Language, 30(5), 513–541. 10.1016/0749-596X(91)90025-F

Kasuga, S., Crevecoeur, F., Cross, K. P., Balalaie, P., & Scott, S. H. (2022). Integration of proprioceptive and visual feedback during online control of reaching. Journal of Neurophysiology, 127(2), 354–372. 10.1152/jn.00639.2020

Krakauer, J. W., Ghez, C., & Ghilardi, M. F. (2005). Adaptation to visuomotor transformations: Consolidation, interference, and forgetting. Journal of Neuroscience, 25(2), 473–478. 10.1523/JNEUROSCI.4218-04.2005

Krakauer, J. W., Hadjiosif, A. M., Xu, J., Wong, A. L., & Haith, A. M. (2019). Motor Learning. In R. Terjung (Ed.), Comprehensive Physiology (1st ed., pp. 613–663). Wiley. 10.1002/cphy.c170043

Leow, L.-A., de Rugy, A., Marinovic, W., Riek, S., & Carroll, T. J. (2016). Savings for visuomotor adaptation require prior history of error, not prior repetition of successful actions. Journal of Neurophysiology, 116(4), 1603–1614. 10.1152/jn.01055.2015

McDougle, S. D., & Taylor, J. A. (2019). Dissociable cognitive strategies for sensorimotor learning. Nature Communications, 10(1). 10.1038/s41467-018-07941-0

Modchalingam, S., Vachon, C. M., Hart, B. M. t., & Henriques, D. Y. P. (2019). The effects of awareness of the perturbation during motor adaptation on hand localization. PLoS ONE, 14(8), 1–20. 10.1371/journal.pone.0220884

Morehead, J. R., Qasim, S. E., Crossley, M. J., & Ivry, R. (2015). Savings upon Re-Aiming in Visuomotor Adaptation. Journal of Neuroscience, 35(42), 14386–14396. 10.1523/jneurosci.1046-15.2015

Neville, K. M., & Cressman, E. K. (2018). The influence of awareness on explicit and implicit contributions to visuomotor adaptation over time. Experimental Brain Research, 236(7), 2047–2059. 10.1007/s00221-018-5282-7

Scott, S. H. (1999). Apparatus for measuring and perturbing shoulder and elbow joint positions and torques during reaching. Journal of Neuroscience Methods, 89(2), 119–127. 10.1016/S0165-0270(99)00053-9

Shadmehr, R., Smith, M. A., & Krakauer, J. W. (2010). Error Correction, Sensory Prediction, and Adaptation in Motor Control. Annual Review of Neuroscience, 33(1), 89–108. 10.1146/annurev-neuro-060909-153135

Singh, K., & Scott, S. H. (2003). A motor learning strategy reflects neural circuitry for limb control. Nature Neuroscience, 6(4), 399–404. 10.1038/nn1026

Taylor, Krakauer, & Ivry. (2014). Explicit and Implicit Contributions to Learning in a Sensorimotor Adaptation Task. Journal of Neuroscience, 34(8), 3023–3032. 10.1523/JNEUROSCI.3619-13.2014

Telgen, S., Parvin, D., & Diedrichsen, J. (2014). Mirror Reversal and Visual Rotation Are Learned and Consolidated via Separate Mechanisms: Recalibrating or Learning *De Novo*? The Journal of Neuroscience, 34(41), 13768–13779. 10.1523/JNEUROSCI.5306-13.2014

Vetter, P., Goodbody, S. J., & Wolpert, D. M. (1999). Evidence for an eye-centered spherical representation of the visuomotor map. Journal of Neurophysiology, 81(2), 935–939. 10.1152/jn.1999.81.2.935

Wang, & Taylor, J. A. (2021). Implicit adaptation to mirror reversal is in the correct coordinate system but the wrong direction. Journal of Neurophysiology, 126(5), 1478–1489. 10.1152/jn.00304.2021

Werner, S., Van Aken, B. C., Hulst, T., Frens, M. A., Van Der Geest, J. N., Strüder, H. K., & Donchin, O. (2015). Awareness of sensorimotor adaptation to visual rotations of different size. PLoS ONE, 10(4), 1–18. 10.1371/journal.pone.0123321

Wilterson, S. A., & Taylor, J. A. (2021). Implicit Visuomotor Adaptation Remains Limited after Several Days of Training. eNeuro, 8(4). 10.1523/ENEURO.0312-20.2021

