## Supplementary Material for "Savings and interference following learning to reach with mirror reversed feedback"

### Supplementary File

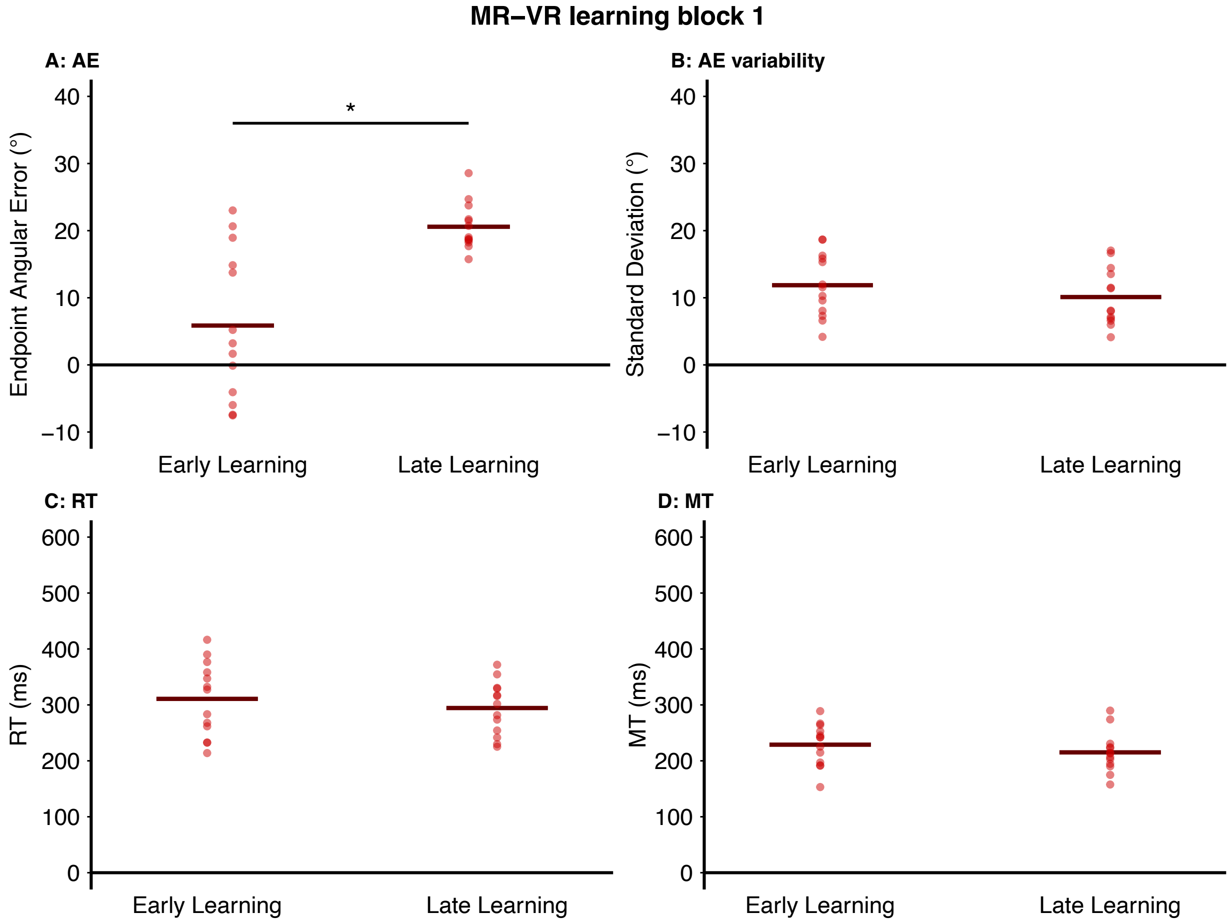


**Figure S1.** Learning to reach with an MR distortion for the MR-VR group, learning block 1. **A:** Endpoint angular error (AE), **B:** AE variability (SD), **C:** reaction time (RT), **D:** movement time (MT), in early and late learning trials in learning block 1. The lines represent the group mean for each variable at a given time point. Circles represent individual participants. Asterisks indicate statistically significant differences between time ($p < 0.05$).
